# Discovery and characterization of highly polymorphic ultra-short STRs for human identification via shotgun sequencing

**DOI:** 10.64898/2026.08.03.742408

**Authors:** Brando Poggiali, Clara I.V. Aagreen, Olivia Luxford Meyer, Alberte Honoré Jepsen, Thorfinn Sand Korneliussen, Marie-Louise Kampmann, Claus Børsting, Jeppe Dyrberg Andersen

## Abstract

Shotgun sequencing (SGS) enables simultaneous interrogation of a broad range of loci across the human genome, even from low-template and highly degraded DNA samples. While human identification traditionally relies on short tandem repeats (STRs) due to their high polymorphism, standard forensic STRs (100-450 bp) are poorly suited for the short read (∼150 bp) constraint of SGS.

The purpose of this study was to evaluate the analysis limitations of standard forensic STRs in SGS data and to identify a novel panel of STRs optimised for short-read genomic data. First, we benchmarked four STR genotyping software tools (STRait Razor, GangSTR, STRinNGS, and HipSTR) by analysing 53 standard forensic STRs in SGS data. HipSTR showed the best performance but achieved only a call rate of 64.5% and an accuracy of 83.8%, and its performance was strongly affected by STR allele length and read depth.

To overcome these constraints, we screened the population-wide 1000 Genomes Project dataset and identified a panel of 265 autosomal ultra-short (< 50 bp) STRs with an effective number of alleles (*A*_*e*_) ranging from 3.0 to 7.5. As few as seven of these loci were sufficient to achieve a Mean Match Probability (MMP) below 1 × 10^−6^. To validate these findings, we developed a custom PCR-based amplicon sequencing panel targeting 97 of the most polymorphic ultra-short STRs and evaluated these in 41 blood samples from Danish individuals. The polymorphic nature of the selected loci was confirmed (*A*_*e*_ranged from 2.4 to 7.2). Our results furthermore demonstrated high concordance between the amplicon panel and SGS-derived genotypes, which substantiates that these ultra-short STRs provide a robust and highly polymorphic alternative for human identification in SGS data.

**Author summary:** Shotgun sequencing (SGS) methods are increasingly being adopted in fields such as forensic genetics. SGS yields large amounts of genetic information by reading short fragments across the entire genome, enabling a wide range of analyses that may be exploited as leads in a police investigation. Human identification has traditionally been based on STR loci with a PCR amplicon length of 100-450 base pairs. However, these loci are often longer than the reads generated by SGS data, which makes them difficult to analyse in a reliable way. In this study, we evaluated four software tools designed to genotype STRs and confirmed the limited ability to genotype traditional forensic STRs in SGS data. To address this limitation, we identified a new set of highly polymorphic ultra-short STRs (less than 50 base pairs in length) that enable robust human identification using SGS data. Despite their shorter length, these loci retain the multi-allelic nature inherent to traditional STRs. This ensures a low random match probability that is comparable with the standard forensic STR panels. The ultra-short STRs may be genotyped from highly degraded DNA and may provide the possibility for complex mixture analysis and multi-donor deconvolution, which makes the STRs uniquely suited for forensic casework.

## 1 Introduction

Short tandem repeats (STRs) are DNA regions consisting of tandemly repeated DNA motifs of 1 to 6 base pairs (bp) [1]. STRs are widespread across the chromosomes (approximately one every 2,000 bp) and represent roughly 3% of the human genome [2]. They are highly polymorphic [3] and may contribute to phenotypic variation [4] and disease development [1]. The polymorphic nature of STRs and ease of detection using length separation methods established them as the marker of choice for human identification (HID) purposes in forensic genetics. Since the early 1990s, STR profiling has been the standard approach for HID [5], and large national DNA databases with thousands or millions of STR profiles are used to identify persons of interest in police investigations on a daily basis [6].

STR profiling is traditionally performed by polymerase chain reaction (PCR), generating amplicons ranging from approximately 100 to 450 bp, depending on the target locus and commercial kit [7,8]. However, forensic samples often contain degraded DNA, and when fragment sizes fall below the PCR amplicon sizes, DNA profiles cannot be obtained [8]. To address this limitation, primer designs were optimised to reduce amplicon length by positioning primers closer to the target repeat region [9]. This strategy led to the development of mini-STRs [10], which demonstrated superior performance in the analysis of highly degraded samples [7,11,12], emphasizing how target-size reduction directly improves DNA profiling success.

Shotgun sequencing (SGS), in which all DNA fragments present in a sample are sequenced, has shown promising results for forensic genetic applications [13,14]. This methodology was extensively used in paleogenetics during the past two decades [15,16], where researchers refined its sensitivity to retrieve genomic information from highly degraded ancient DNA [17]. Several forensic sample types, including skeletal remains and burned bodies, exhibit DNA degradation patterns that are similar to those of ancient DNA samples, suggesting that SGS could yield informative genetic data when conventional techniques fail [18–20]. In addition, SGS generates reads that cover the whole genome and provides the opportunity to conduct multiple genetic analyses (e.g., HID, investigative genetic genealogy, and biogeographic ancestry inference) simultaneously [13,21,22]. Nevertheless, SGS typically generates DNA reads of 30-150 bp, which may be too short for genotyping some STRs since the reads should span the entire repeat region and some parts of the flanking sequences on both sides of the STR [1,23]. Standard STRs used for HID were not historically selected to fit this constraint, and therefore, the current standard STRs used for HID may be too long for SGS [13]. STRs with shorter allele lengths and flanks with little variation may be more optimal for SGS. Moreover, highly degraded forensic samples often contain DNA with fragment sizes significantly shorter than 150 bp. Burned bodies, compromised bones, touch DNA, and rootless hair shafts are enriched in DNA fragments in the range of ∼35 to 90 bp [19,24–27], which often makes standard STR profiling unfeasible. SNP markers have also been considered for HID [28–30]. Their reduced DNA length (single base) makes them particularly suitable for degraded samples. However, their low polymorphism results in lower discriminatory power compared to STRs and makes them less suitable for the interpretation of DNA mixture samples, which are often encountered in forensic casework [31].

In this study, we aimed to assess the performance of STR genotyping software tools on traditional STRs in SGS data and to identify a novel panel of STRs optimised for HID in short-read genomic data. First, we benchmarked four STR genotyping software tools (STRait Razor [32], STRinNGS [23], GangSTR [33], and HipSTR [34]) by genotyping 53 standard forensic STR loci in SGS data with known STR profiles. Next, we used HipSTR to identify polymorphic (*A*_*e*_> 3), ultra-short (< 50 bp) STR loci in the large-scale 1000 Genomes Project (1KGP) data. Finally, we developed a custom PCR-based amplicon sequencing panel targeting the candidate STRs, tested it on both high-quality and degraded DNA samples, and characterised the allele frequencies in a Danish cohort.

## 2 Results

### 2.1 Benchmarking of STR genotyping software tools using shotgun sequencing data

We assessed the performance of four STR genotyping software tools: STRait Razor, GangSTR, STRinNGS, and HipSTR, for genotyping 53 forensic STRs (from the Verogen ForenSeq MainstAY kit, hereafter MainstAY) across 12 in-house SGS samples with known STR profiles. These 12 SGS samples originated from three healthy individuals sequenced using a combination of two extraction kits and two library preparation kits (see Materials and Methods). For each tool, detected genotypes were classified as correct calls, drop-ins, drop-outs, or no calls (Supplementary Figure S1). When no sequencing depth (DP) threshold was applied, HipSTR exhibited the highest percentage of correct calls (58.2%), while STRinNGS showed the lowest (15.6%) (Supplementary Figure S1A). Drop-in and drop-out events were more frequent for STRait Razor and GangSTR than for STRinNGS and HipSTR. Applying a DP threshold (eight for autosomal STRs and four for Y-STRs) reduced the percentage of correct calls, drop-ins, and drop-outs, while increasing the proportion of no calls for all four software tools (Supplementary Figure S1B). HipSTR maintained the highest percentage of correct calls (54.1%), while STRinNGS and STRait Razor had the lowest number of drop-outs and drop-ins.

We next evaluated call rate and accuracy conditional on a genotype call, both with and without DP filtering (Figure 1). Without a DP threshold, call rates ranged from 27.6% for STRinNGS to 95.7% for GangSTR. Applying the DP threshold reduced call rates across all software tools, most notably for STRait Razor, which decreased from 76.0% to 30.9%. Accuracy conditional on a genotype call varied across the software tools. Without DP filtering, HipSTR showed the highest accuracy (79.4%), whereas GangSTR showed the lowest (46.0%). With a DP threshold, the accuracy increased for all software tools, with HipSTR reaching the highest value (83.8%) and GangSTR remaining the lowest (55.0%). Overall, HipSTR consistently showed higher accuracy while maintaining a comparatively high call rate.

**Figure 1.**
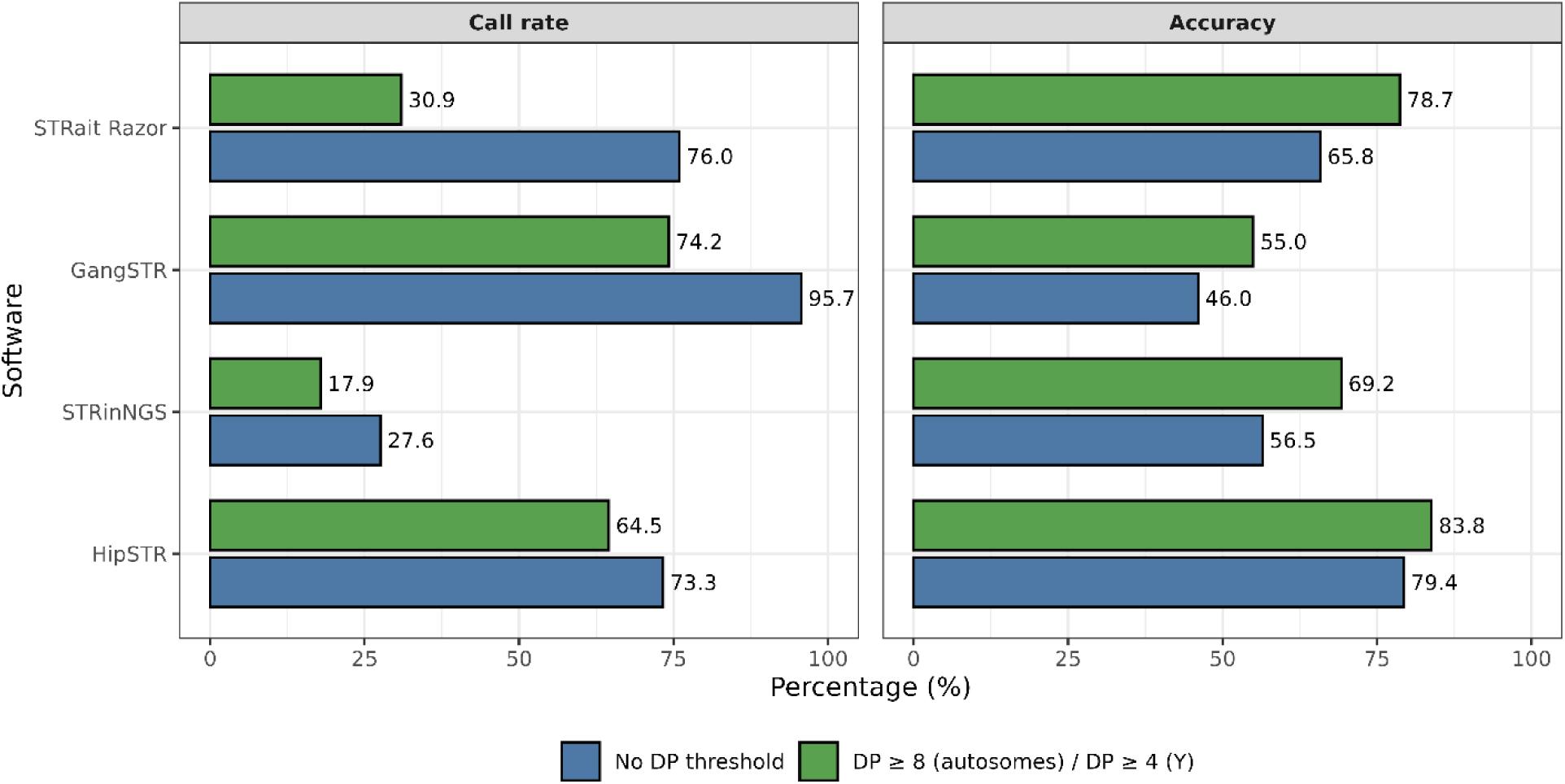
Call rate and accuracy of four STR genotyping software tools on 53 forensic STRs in shotgun sequencing data. Plots show call rate and accuracy (%) for each tool, run on the 12 in-house shotgun sequencing samples under two sequencing depth (DP) conditions: no threshold and a minimum depth of eight reads.

Locus-level performance for HipSTR is reported in Supplementary Table S1. The STR loci D16S539, D22S1045, D2S1338, D8S1179, DYS439, DYS522, and DYS576 were genotyped correctly by HipSTR in 100% of the samples, while four additional loci (D2S441, DYS437, DYS533, and DYS19) were either correctly called or not called across all samples. In contrast, seven STR loci (D21S11, DYF387S1, DYS385, DYS389II, DYS448, DYS612, and SE33) were not called in any of the 12 samples.

We further investigated the effect of STR allele length and DP on genotyping performance. Both call rate and accuracy decreased with increasing STR allele length (Supplementary Figure S2). For alleles longer than 100 bp, call rates approached zero for most software tools, whereas GangSTR continued to report genotype calls at these lengths; however, these calls had low accuracy. Analysis of genotyping performance as a function of DP revealed a consistent pattern across all four software tools (Supplementary Figure S3): DP had little impact on call rate, but accuracy was low at low DP values and increased proportionally with sequencing depth. No DP threshold was set in the genotyping software tools, which therefore always called a genotype.

### 2.2 Identification of highly polymorphic ultra-short STRs

To identify STRs that may be of use for HID in SGS data, we screened the HipSTR reference catalogue containing 1,638,945 human STR loci. This catalogue included 840,248 mononucleotide STRs, 300,374 dinucleotide STRs, 81,177 trinucleotide STRs, 241,115 tetranucleotide STRs, 101,982 pentanucleotide STRs, and 74,049 hexanucleotide STRs, with reference sequence lengths ranging from 9 to 132,210 bp. The top 100 most common repeat motifs are listed in Supplementary Table S2. The catalogue was filtered as illustrated in Figure 2. First, loci were retained based on repeat unit length (> 2 bp), reference sequence length (15-50 bp), and non-overlap with gene exons, yielding 382,410 STR loci.

**Figure 2:**
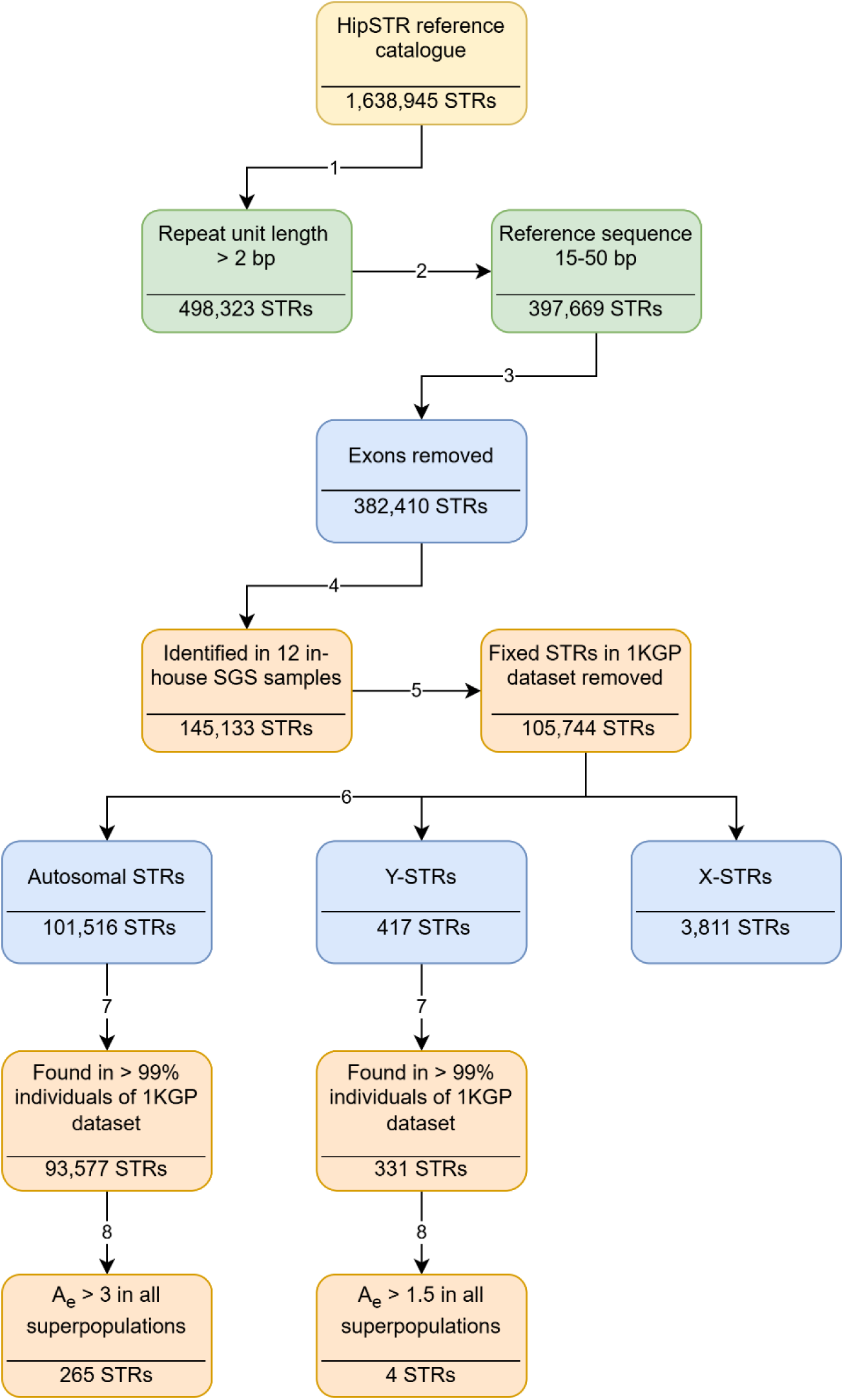
Flowchart showing the workflow for the discovery of highly polymorphic ultra-short STRs. The workflow includes filtering based on STR length (green), genotypes derived from sequencing data (orange), and STR genomic location (blue). STRs: short tandem repeats; bp: base pair; SGS: shotgun sequencing, 1KGP: 1000 Genomes Project, *A*_*e*_: effective number of alleles, Y-STRs: Y-chromosomal STRs, X-STRs: X-chromosomal STRs. Figure created using draw.io

These loci were genotyped in the 12 in-house SGS samples using HipSTR to ensure that the STRs could be detected in SGS data. Only loci that had concordant genotypes across all samples from the same donor were retained, which reduced the number of candidate STR loci to 145,133.

These loci were subsequently genotyped in 2,504 unrelated individuals from five superpopulations of the 1KGP dataset, and the number of alleles per locus was calculated. The distribution of STR loci by the number of alleles is shown in Supplementary Figure S4A. We identified 39,389 loci with a fixed allele across all superpopulations, which were excluded from further analysis as non-polymorphic. We also observed that STRs with shorter motifs tended to exhibit a higher number of alleles (Supplementary Figure S4B). Then, we classified the STRs as autosomal STRs (aSTRs), Y-chromosomal STRs (Y-STRs), and X-chromosomal STRs (X-STRs). We excluded X-STRs from further downstream analysis, as they are suboptimal for HID purposes, and only retained aSTR and Y-STR loci genotyped in at least 99% of the individuals in the 1KGP dataset to ensure a high call rate. This reduced the number to 93,577 aSTRs and 331 Y-STRs.

To identify the most polymorphic STRs suitable for HID, allele frequencies were calculated for each locus, and *A*_*e*_values were estimated per superpopulation for aSTRs and Y-STRs (Supplementary Figures S5 and S6). Most aSTRs and Y-STRs had *A*_*e*_ values close to 1, indicating low degrees of polymorphism. To focus on the most informative STRs for HID, we selected aSTRs and Y-STRs with minimum *A*_*e*_values of 3 and 1.5 in all five superpopulations, respectively. This filtering resulted in 265 aSTRs and four Y-STRs, which we defined as relevant candidate STRs (Supplementary Tables S3 and S4). The 265 aSTRs were mainly tri- and tetranucleotides (Supplementary Figure S7A), with the top four repeat motifs being AAT, AAAT, ATTT, and ATT (Supplementary Figure S7B). Only two forensic STR loci from the MainstAY kit were identified as highly polymorphic ultra-short STRs: D16S539 and DYS522.

### 2.3 Mean match probability of sets of independent highly polymorphic ultra-short STRs

To calculate the discriminatory power of the highly polymorphic ultra-short STRs for HID, we selected independent STR loci and calculated the mean match probability (MMP) (see Materials and Methods). From the 265 aSTRs identified in the previous analysis, we selected one locus per chromosome arm in the GRCh38 reference genome. Because some short arms are absent or not represented in GRCh38, this resulted in a final set of 39 loci. MMP values were calculated for each of the five superpopulations using allele frequencies derived from 1KGP genotypes (2,504 individuals). The lowest MMP (i.e., highest discriminatory power) was observed in the African superpopulation (1.46 x 10^-50^) and the highest in the East Asian superpopulation (3.44 x 10^-44^) (Figure 3B).

**Figure 3:**
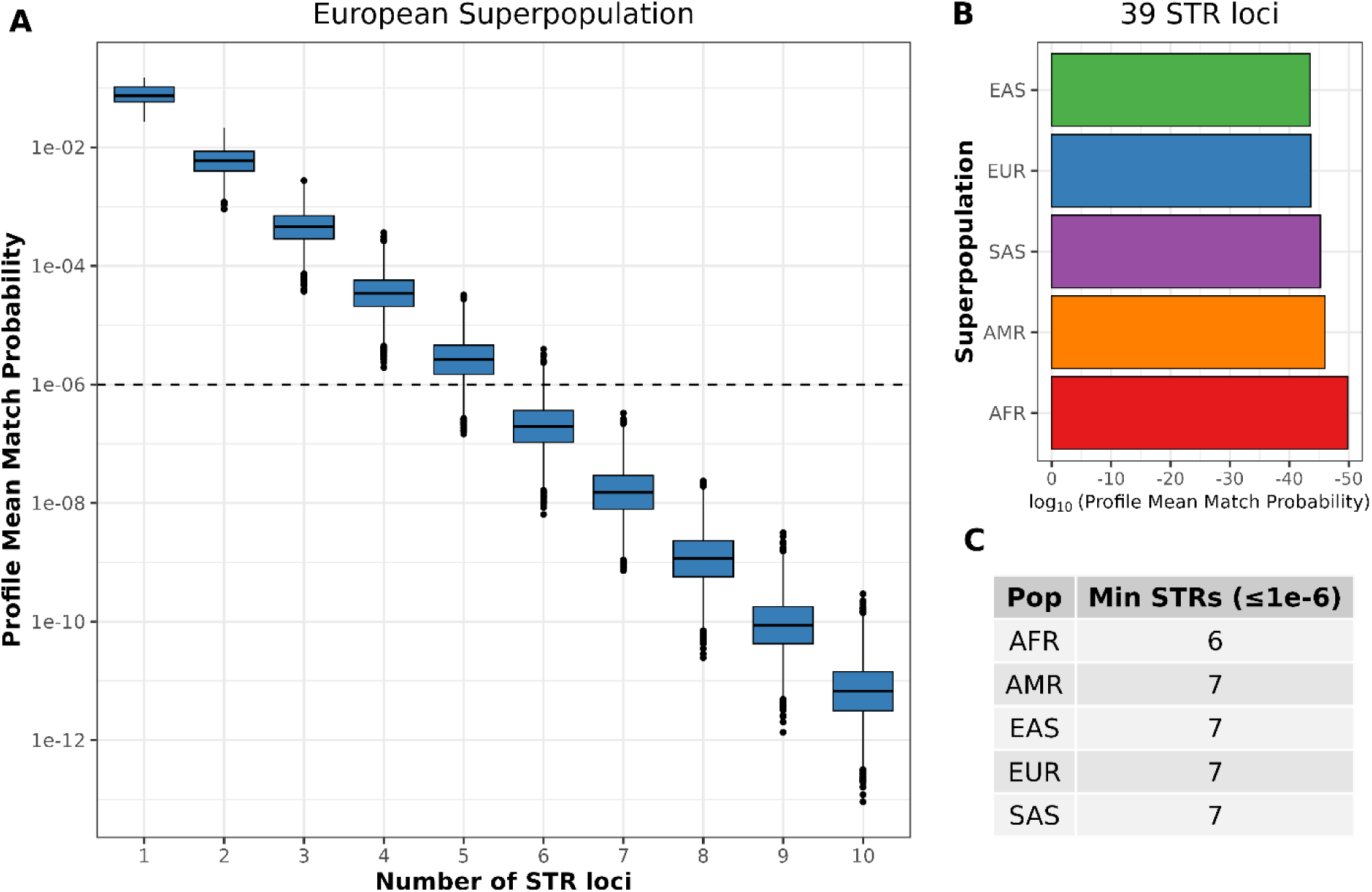
Mean match probabilities (MMPs) based on 39 autosomal STR loci from the list of highly polymorphic ultra-short STR loci. **A**) MMP of 10,000 random combinations of STRs ranging from one to 10 loci in the European superpopulation. The dotted line indicates the MMP threshold of a 1 × 10^−6^. **B**) MMP of the 39 STR loci across the five superpopulations: African (AFR), European (EUR), American (AMR), and East (EAS) and South Asian (SAS). **C**) Minimum number of STR loci to reach an MMP lower than 1 × 10^−6^.

In practice, forensic trace samples may yield incomplete STR profiles due to low DNA quantity and/or degradation. To evaluate the impact of missing loci in the final set of 39 loci, we simulated 10,000 random combinations of reduced STR sets ranging from one to ten loci and calculated the corresponding MMPs for each superpopulation (Figure 3A and Supplementary Figure S8). To consistently achieve an MMP below 1 × 10^−6^ — the symbolic threshold used for HID — a minimum of six loci was sufficient for the African superpopulation, while at least seven loci were required for all other superpopulations (Figure 3C).

### 2.4 Targeted sequencing of highly polymorphic ultra-short STRs in a Danish cohort

To validate the polymorphic nature of the identified STR loci, we developed a custom PCR-based amplicon sequencing panel targeting 97 of the most polymorphic STRs (highest *A*_*e*_) (Supplementary Table S5). The PCR product input sizes ranged from 70 to 102 bp (mean 87 bp). The actual amplicon sizes including primers ranged from 126 to 140 bp (mean 138 bp). Using this panel, we performed targeted sequencing of blood samples from 41 donors to characterise allele frequencies in a Danish population. In addition, four telogen hair samples were sequenced to evaluate the panel’s performance on low-template and fragmented DNA. Because we aimed to characterise length-based and sequence-level STR genotypes, together with flanking-region variability, from targeted amplicon sequencing data, we used STRinNGS, which supports all of these analyses.

The call rate for the blood samples exceeded 91.8% in all individuals, whereas the four hair samples showed a substantially lower call rate (2.1-17.5%), indicating poor STR genotyping performance in highly degraded samples (Supplementary Figure S9). Nevertheless, we observed genotype concordance at 21 STR loci between hair and blood samples, while five showed one allele mismatch (Supplementary Table S6). We then assessed the individual STR call rate, excluding the hair samples. Most STR loci showed a call rate of 100%, while four loci failed to generate genotypes in any of the samples, and five loci suffered from locus drop-outs in a few samples (Supplementary Table S7). We then assessed genotype concordance between the three individuals sequenced with SGS and the amplicon panel across 93 STR loci (Supplementary Table S8). A total of 90, 89, and 91 concordant genotypes were observed for donors A, B, and C, respectively, indicating high concordance between the two technologies. Discordant genotypes were identified at two STR loci, Human_STR_200966 and ATCT053. Of the 93 STRs with genotyping data, 67 were simple STRs, 20 were compound STRs, and six were complex STRs. A total of 16 SNPs and one indel were observed in the flanking regions, and this variation was also detected and added to the STR allele nomenclature. The allele frequencies and proposed allele nomenclature for the 93 STRs are shown in Supplementary Table S9.

Finally, *A*_*e*_values were calculated from the 41 Danish donors and compared with those derived from the 1KGP (Figure 4A and Supplementary Table S10). Similar *A*_*e*_ values were observed between the two datasets, although some differences were present. Because STRinNGS reports both length-based and sequence-level alleles, including SNPs in the flanking regions, we estimated the *A*_*e*_using both allele definitions. As expected, sequence-level alleles had higher *A*_*e*_values than length-based alleles for 36 of the sequenced STRs (Figure 4B and Supplementary Table S10).

**Figure 4.**
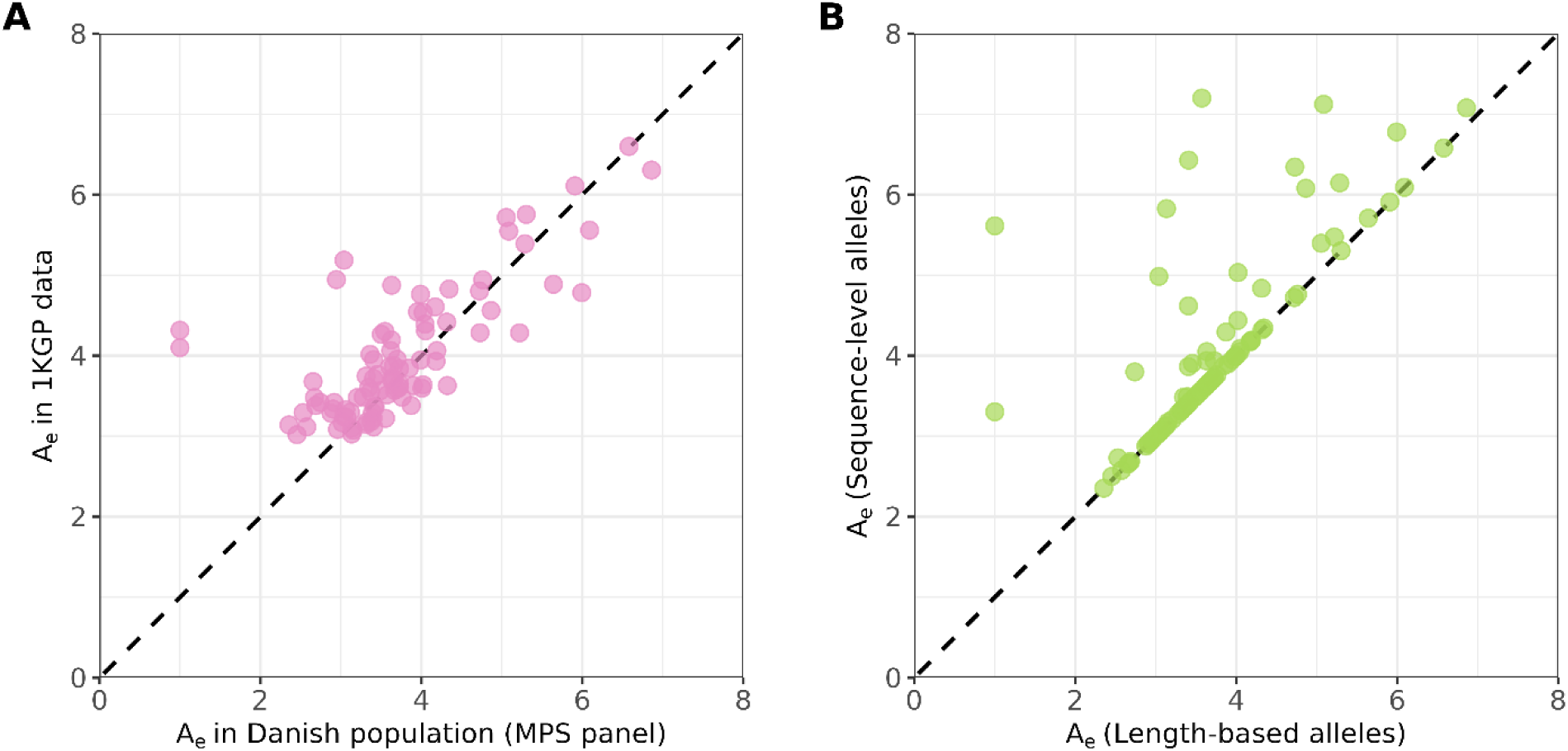
Effective number of allele (*A*_*e*_) values obtained from the 41 Danish donors sequenced with the custom STR panel. **A)** Comparison of *A*_*e*_ values calculated in the European superpopulation of 1000 Genomes Project (1KGP) and the 41 Danish donors. **B)** *A*_*e*_ values calculated using length-based and sequence-level alleles in blood samples of 41 Danish donors.

## 3 Discussion

The cost of generating a human genome has declined drastically over the past two decades [35]. As a result, high-throughput sequencing has become increasingly accessible, and SGS is slowly being introduced into forensic genetic laboratories worldwide [14,21,22]. SGS combines the ability to simultaneously interrogate all loci of the human genome with the sensitivity required for low-template DNA samples [20]. However, with the typical short-read length of 150 bp imposed by SGS, the standard forensic STRs used for HID are not suitable. This was also observed when we investigated the standard STRs using the four STR genotyping software tools: STRait Razor, STRinNGS, GangSTR, and HipSTR. Indeed, sequencing-based STR genotyping requires reads to span the full STR region and the flanking regions. This is particularly complicated in highly degraded DNA samples, where DNA fragments are often < 100 bp [19,24–27]. To overcome this limitation, we identified highly polymorphic ultra-short STRs with genomic lengths < 50 bp that enable HID with SGS and PCR-based targeted sequencing approaches.

STR genotype calling relies on specialised bioinformatic software tools. We benchmarked four STR genotyping software: STRait Razor, STRinNGS, GangSTR, and HipSTR. STRait Razor and STRinNGS were specifically developed for forensic (PCR amplicon–based) applications, whereas GangSTR and HipSTR are general-purpose STR genotyping software tools. These software tools were selected because they are freely available, well-maintained, easy to deploy locally, and have shown high genotyping capabilities in previous studies. However, they have not been jointly benchmarked in a single study [32,36–38]. HipSTR showed the best performance, achieving a call rate of 64.5% and an accuracy of 83.8% across the 53 forensic STRs included in the MainstAY kit. These results are partially aligned with those of Valle-Silva *et al.* [39], who also reported higher call rates for HipSTR compared to STRait Razor, although not higher accuracy. Interestingly, they reported higher performance (call rate of 84.9% and accuracy of 94.01%) for HipSTR than that observed in our study. This discrepancy can be attributed to different methodological approaches between the two studies. First, their benchmark focused exclusively on aSTRs, whereas our analysis also included Y-chromosomal STRs, which we found to have lower genotyping performance. Second, they used a targeted sequencing panel rather than SGS, thereby overcoming the 150 bp read length constraint of this technology. Valle-Silva *et al.* [39] also observed that the tool toaSTR slightly outperformed both HipSTR and STRait Razor in terms of call rate and accuracy. However, toaSTR was excluded from our benchmarking because it requires the upload of raw sequencing data to an external server [40], which raises concerns for data privacy and regulatory limitations. Recently, a new tool, STRsensor, was released and showed improved performance over HipSTR in simulated whole-genome sequencing data [41]. Future studies should compare the performance of this tool for the analysis of standard forensic STR loci. Nevertheless, HipSTR has been shown to be a robust, fast, scalable, and flexible genotyping tool [34,42], and based on both our results and previous results [39], it was selected for genotyping the highly polymorphic ultra-short STRs in 2,504 unrelated individuals of the 1KGP. Moreover, the recent release of a graphical interface (HipSTR-UI) facilitates the adoption of this tool by forensic practitioners with non-bioinformatic experience [43].

Locus-specific performance assessed with HipSTR revealed high variability across the 53 standard forensic STRs. Some loci were consistently genotyped correctly in all SGS samples, whereas others systematically failed. In several cases, genotyping performance could be explained by STR length. For example, D21S11, with a reference core sequence length of 127 bp, showed a call rate of 0%, consistent with previous observations [42]. When we systematically assessed allele length versus genotyping accuracy, we observed that alleles longer than ∼100 bp were not correctly genotyped. Longer alleles have a lower probability of being fully spanned by reads, including their flanking regions. Our result is consistent with previous observations that shorter STR alleles yield better genotyping performance [44]. However, STR length alone did not fully explain performance variability, indicating that additional factors are involved. Erroneous genotype calls could also be induced by stutter artefacts generated during PCR amplification. Factors such as repeat number, motif type (e.g., di- or trinucleotide), sequence composition (i.e., AT content), repeat structure complexity, and flanking regions influence stutter generation [45–49]. These factors should therefore be considered during marker selection. Furthermore, we observed that genotyping accuracy was directly correlated with DP, highlighting the importance of defining a locus-specific minimum DP to balance call rate and accuracy.

In summary, sequencing-based STR genotyping performance is determined by a complex interplay among read length, STR intrinsic properties, DP, and the bioinformatic tool used for genotyping.

To identify STRs compatible with SGS, we screened 1,638,945 loci from the HipSTR catalogue and selected 265 highly polymorphic ultra-short aSTRs and four Y-STRs suitable for HID. We applied stringent filtering criteria to retain loci optimised for accurate genotyping. Mono- and dinucleotide STRs were excluded due to their propensity to generate stutter artefacts [50–52]. We also restricted the analysis to loci with a reference length between 15 and 50 bp. Previous work showed that the reference length approximates the median allele length distribution [53]; thus, shorter loci (<15 bp) were expected to exhibit limited polymorphism. In contrast, loci > 50 bp were more likely to exceed the read length constraint of SGS and degraded DNA. To further ensure robustness, we retained loci that were consistently genotyped across the same individuals in our SGS data.

Allele frequencies and effective number of alleles (A_e_) across the human population were estimated using the 1KGP dataset (2,504 individuals from 26 populations). We selected highly polymorphic loci, resulting in 265 aSTRs (A_e_ range 3-7.5) and four Y-STRs (A_e_ range 2.23-2.82).

Notably, the A_e_ distribution overlaps with that of standard forensic STRs [54]. Among the 53 STRs in the MainstAY kit, only D16S539 and DYS522 passed all selection criteria, highlighting the limited compatibility of standard forensic STRs with SGS constraints. As few as seven of the identified aSTRs were sufficient to achieve an MMP below the symbolic threshold of 1 × 10^−6^. Given that forensic samples often yield incomplete profiles, the availability of multiple highly informative STRs increases the likelihood of successful identification.

We further developed a custom multiplex amplicon panel targeting 97 of the most polymorphic aSTRs and validated their genetic variation in 41 individuals from the Danish population. High concordance was observed between genotypes obtained from blood samples using the multiplex panel and SGS data, with only two loci (Human_STR_200966 and ATCT053) showing discordance. This result supports the robustness of both sequencing approaches and confirm the suitability of the selected STRs for HID with SGS. The A_e_ values observed in the Danish cohort were consistent with those derived from the global population, supporting the effective polymorphic nature of the loci. Additionally, incorporating sequence-level allelic variation increased the A_e_ and, consequently, discriminatory power.

The panel performance was substantially reduced in hair samples. This result was expected because the final amplicon size ranged from 126 to 140 bp and hair samples typically contain DNA fragment size lower than 100 bp [24,25]. Nevertheless, 21 out of 26 genotypes were concordant with those obtained from blood, demonstrating partial robustness under challenging conditions.

Although STRs have been historically chosen for HID, alternative genetic markers, such as SNPs and microhaplotypes, have also been previously explored [55,56]. SNPs are optimal for degraded samples due to their short length, but their limited polymorphism necessitates a larger number of markers and complicates mixture interpretation [31]. Microhaplotypes consist of closely linked SNPs within short genomic region (< 300 bp) and can achieve the level of polymorphism of forensic STRs [54]. Previous studies have developed microhaplotypes with length below 50-70 bp; although their A_e_ values were lower than those of standard STRs, they achieved high discriminatory power with relatively few markers [57,58]. A major advantage of microhaplotypes over STRs is the absence of stutter artefacts, which facilitate mixture interpretation [55,57]. Moreover, microhaplotypes do not exhibit allele length imbalance, as observed in STRs, where shorter alleles are preferentially amplified and genotyped, particularly in degraded samples [55]. However, microhaplotypes have been typed with targeted amplicon assays [55], and their performance in SGS data has not yet been explored. In addition, the bioinformatic ecosystem for microhaplotype analysis is still limited, although rapidly expanding [59–62]. Future studies should investigate their integration into an SGS-based forensic workflow.

In conclusion, we identified a panel of highly polymorphic ultra-short STRs that are suitable for individual genetic discrimination. These markers are particularly beneficial when SGS is applied to both high-quality and degraded DNA samples. For genotyping these STRs in SGS data, we recommend the use of the bioinformatic tool HipSTR.

## 4 Materials and methods

### 4.1 Ethical approval

The study was approved by the Scientific Ethics Committees for the Capital Region of Denmark (H-22034131). Written informed consent was collected from all participants. Participants were recruited between 02 December 2022 and 10 June 2024. The samples are stored in pseudonymised form in a biobank that is registered at the University of Copenhagen’s joint records of processing of personal data in research projects and biobanks (514–0789/23–3000) and complies with the rules of the General Data Protection Regulation (Regulation (EU) 2016/679).

### 4.2 Benchmarking of STR genotyping software tools in shotgun sequencing data

#### 4.2.1 Shotgun sequencing

SGS data was previously generated by Kampmann *et al.* [13]. A total of 12 samples were included in this study (Supplementary Table S11). Briefly, buccal swabs from three healthy Danish donors were extracted with the EZ1&2 DNA Investigator Kit (Qiagen, Hilden, Germany). Libraries were prepared using either the NEBNext® Ultra™ II DNA Library Prep Kit for Illumina (New England Biolabs, Ipswich, MA, USA) or the TruSeq Nano DNA Low Throughput Library (Illumina, San Diego, CA, USA). All the DNA extracts were fragmented using an S220 Focused-ultrasonicator (Covaris) following the settings from the 350 bp TruSeq Nano DNA Low Throughput Library protocol, and the final libraries were sequenced on a NovaSeq 6000 Illumina sequencing system using 150 bp paired-end sequencing. Sequence data analysis was described in detail by Kampmann *et al.* [13]. The genome-wide coverage ranged from 9.3× to 44.2×.

#### 4.2.2 Bioinformatic analysis and software tools comparison

The genotyping performance of the four STR genotyping software tools (STRait Razor v3.0 [32], GangSTR v2.5.0 [33], STRinNGS v2.1 [23], and HipSTR v0.6.2 [34]) were tested on SGS samples with known STR profiles. These software tools were selected because they are freely available, well-maintained, easy to install, and showed high genotyping performance in previous studies [32,36–38]. A detailed description of the software tools setting, input files, and command lines used for each tool is provided in Supplementary Materials. All four software tools were applied to the 53 forensic STR loci included in the MainstAY kit (Verogen, San Diego, CA, USA) [63], which comprises 28 aSTRs, 25 Y-STRs, and the marker Amelogenin for sex determination. Detailed information on the genomic coordinates, repeat unit, and locus length of the MainstAY STRs is provided in Supplementary Table S12.

The STR genotypes generated by the four software tools were compared with reference STR profiles generated using the MainstAY kit (Verogen, San Diego, CA, USA) and capillary electrophoresis (3500xL Genetic Analyzer, Thermo Fisher Scientific, Waltham, MA, USA), which is considered the gold standard for forensic STR analysis. For each STR genotyping tool, the proportion of correct calls, drop-ins, drop-outs, and no calls were calculated. The call rate was defined as the number of loci with a genotype call (correct calls, drop-ins, or drop-outs) divided by the total number of loci analysed. A call was considered correct when the length-based genotype matched the reference, a drop-in when an incorrect allele was present in the genotype, and a drop-out when one of the two alleles failed to be detected. Accuracy was calculated as the number of correct calls divided by the total number of genotype calls (correct calls, drop-ins, and drop-outs) × 100. All plots were created with R (version 4.4.1) [64] and ggplot2 (version 3.5.1) [65].

### 4.3 Discovery of highly polymorphic ultra-short STRs

#### 4.3.1 1000 Genomes Project (1KGP) data

For the identification of highly polymorphic ultra-short STRs, data from 1KGP phase 3 were used. Specifically, 2,504 CRAM files from 2,504 unrelated individuals (1,271 females and 1,233 males) representing five superpopulations – East Asian, European, African, American, and South Asian – were downloaded from ftp://ftp.sra.ebi.ac.uk/vol1/run/ERR324/ and ftp://ftp.sra.ebi.ac.uk/vol1/run/ERR323/ [66]. Additional information on the superpopulations can be found in Supplementary Table S13. The data was sequenced to approximately 30× coverage and aligned to the human reference genome GRCh38 by the New York Genome Center [67].

#### 4.3.2 Pipeline for the discovery of highly polymorphic ultra-short STRs

The genome-wide STR reference catalogue “hg38.hipstr_reference.bed.gz” was downloaded from the HipSTR reference page [68]. This file contains 1,638,945 STR loci with repeat unit lengths ranging from 1 to 6 bp. To discover highly polymorphic ultra-short STRs suitable for HID in highly degraded samples and short-read sequencing, the reference file was filtered according to the steps outlined below. An overview of the process is shown in Figure 2.

1. To reduce genotyping errors due to PCR stutter, STRs with repeat motifs ≤2 bp were excluded from the catalogue.
2. STR loci with reference sequence < 15 bp or > 50 bp were removed. It was assumed that STR loci with reference lengths < 15 bp would have limited allelic variation, as observed by Shi *et al.* [53], whereas loci > 50 bp would be less suitable for HID in degraded samples.
3. STR loci located within gene exons (minimum overlap of 1 bp) were discarded to avoid inclusion of loci that may reveal phenotypic information. This filtering step was performed using BEDTools subtract (version 2.22.1) [69]. Exon coordinates for the human genome (Ensembl release 112) were obtained in BED format using Ensembl’s Biomart [70,71].
4. To identify STR loci that were detectable in SGS data, HipSTR was applied to the 12 in-house SGS samples using the filtered STR catalogue from the previous step. STR loci with identical genotypes across all samples originating from the same donor were retained.
5. The resulting STR reference catalogue was used to genotype STRs in 2,504 individuals from the 1KGP using HipSTR. STR loci that were fixed (i.e., showing the same allele in all individuals) were discarded.
6. STRs were divided into aSTRs, Y-STRs, and X-STRs.
7. To identify STR loci with high genotyping call rates, only loci detected in at least 99% of the 1KGP individuals were retained.
8. To estimate the polymorphic nature of these STR loci, allele frequencies (*p*_*i*_) were calculated for each locus and used to estimate the *A*_*e*_:

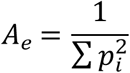 Finally, STR loci with an *A*_*e*_ larger than 3 for aSTRs and larger than 1.5 for Y-STRs were retained. No filtering was performed on the X-STRs.

### 4.4 Mean match probability of the highly polymorphic ultra-short STRs

To calculate the MMP, we selected STR loci that were not in linkage disequilibrium by retaining only loci located on different chromosome arms (p or q chromosome arm), based on centromere coordinates (GRCh38 assembly) downloaded from the UCSC Table Browser [72]. For each chromosome arm, the STR locus with the highest *A*_*e*_value was retained. The MMP of these STRs was calculated from allele frequencies estimated using the 1KGP genotypes as described in [35]. First, the locus-specific MP was calculated for each STR locus by summing the squared frequency of each possible genotype, assuming Hardy-Weinberg equilibrium. For example, at a locus with three alleles: A, B, and C with frequencies a, b, and c, respectively, the locus-specific match probability was:

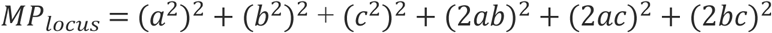

Then, the MMP across all STR loci is calculated as the product of the locus-specific MP values. To determine the minimum number of the highly polymorphic ultra-short STRs required for HID, 10,000 random combinations of loci were generated for each possible set size ranging from 1 to 10 loci. For each set size, the MMP was computed across all random combinations, and the resulting distributions were visualised as boxplots.

### 4.5 Development of a custom PCR-based amplicon sequencing panel targeting the highly polymorphic ultra-short STRs

The DesignStudio Assay Design Tool (Illumina, San Diego, CA, USA) [73] was used to design a custom AmpliSeq panel with 103 out of the 265 most polymorphic aSTR loci. The targeted positions were extended by ±2 bp from the reported STR start and end coordinates. Six of the 103 target STRs were not included in the final AmpliSeq design because primers for these positions were deemed “undesignable”. The final panel included 97 STRs and 10 sex determination markers (Amelogenin and 9 SNPs) (Supplementary Table S5).

### 4.6 Characterization of the highly polymorphic ultra-short STR panel in blood and hair samples in a Danish cohort

Whole blood samples from 41 healthy Danish individuals were collected in EDTA whole blood collection tubes. Four telogen hair samples from two healthy Danish individuals were collected. Two of them were 9 cm long, and the other two were 2 cm long. Genomic DNA extraction was carried out from blood samples using the QIAamp DNA Blood Mini Kit (Qiagen, Hilden, Germany) according to the manufacturer’s guidelines. Extracted genomic DNA was quantified with the Quantifiler™ Trio DNA Quantification kit (Thermo Fisher Scientific, Waltham, MA, USA) using the Applied Biosystems™ 7500 real-time PCR system (Thermo Fisher Scientific, Waltham, MA, USA) according to the manufacturer’s protocol. The hairs were washed using Terg-a-zyme as described by Brandhagen *et al*. [24]. DNA was extracted using the buffer described by Pedersen *et al*. [74], followed by purification using High Pure Viral Nucleic Acid Large Volume Kit (Roche Diagnostics, Basel, Switzerland). The concentration of the extracted DNA was determined using the Quantifiler™ Trio DNA Quantification Kit.

Sequencing libraries were built using the AmpliSeq™ Library Prep Kit following the manufacturer’s instructions and sequenced on the MiSeq System (Illumina, San Diego, CA, USA). The MiSeq output, in the form of binary base call (BCL) files, was converted to FASTQ files with bcl2fastq2 (Illumina, San Diego, CA, USA). FASTQ files were used as input for STRinNGS [23], along with the reference genome GRCh38 and a custom INI file containing STR locus coordinates, motifs, SNPs in flanking regions, and locus analysis parameters (Supplementary File 2). The repeat motif, as well as start and stop positions, were adjusted for some of the STRs to include all allelic variations and account for PCR primer positions. These modifications generated discrepancies in locus definitions and genotyping conventions between the amplicon panel and SGS approaches. To enable direct comparison of the STR genotypes obtained with the amplicon panel (genotyped using STRinNGS) and SGS data (genotyped using HipSTR), allele lengths were harmonised across the two genotyping approaches. Locus-specific allele-length corrections were applied and reported in Supplementary Table S14.

## CRediT

- Brando Poggiali: Conceptualization, Methodology, Formal analysis, Investigation, Supervision, Data Curation, Writing – Original Draft, Visualization
- Clara I.V. Aagreen: Methodology, Software, Formal analysis, Investigation, Data Curation, Visualization, Writing – Review & Editing
- Olivia Luxford Meyer: Methodology, Software, Formal analysis, Writing – Review & Editing
- Alberte Honoré Jepsen: Methodology, Writing – Review & Editing
- Marie-Louise Kampmann: Conceptualization, Supervision, Writing – Review & Editing
- Thorfinn Sand Korneliussen: Supervision, Writing – Review & Editing
- Claus Børsting: Conceptualization, Methodology, Formal analysis, Supervision, Writing – Review & Editing
- Jeppe Dyrberg Andersen: Conceptualization, Investigation, Supervision, Writing – Review & Editing

## References

1. Rajan-Babu IS, Dolzhenko E, Eberle MA, Friedman JM. Sequence composition changes in short tandem repeats: heterogeneity, detection, mechanisms and clinical implications. Nat Rev Genet. 2024 Jul 11;25(7):476–99. doi:10.1038/s41576-024-00696-z

2. Fan H, Chu JY. A Brief Review of Short Tandem Repeat Mutation. Genomics Proteomics Bioinformatics. 2007 Mar 1;5(1):7–14. doi:10.1016/S1672-0229(07)60009-6

3. Willems T, Gymrek M, Highnam G, Mittelman D, Erlich Y. The landscape of human STR variation. Genome Res. 2014 Nov;24(11):1894–904. doi:10.1101/gr.177774.114

4. Lamkin M, Gymrek M. The emerging role of tandem repeats in complex traits. Nat Rev Genet. 2024 Jul 7;25(7):452–3. doi:10.1038/s41576-024-00736-8

5. Roewer L. DNA fingerprinting in forensics: past, present, future. Investig Genet. 2013;4(1):22. doi:10.1186/2041-2223-4-22

6. Wickenheiser RA. Expanding DNA database effectiveness. Forensic Sci Int. 2022;4:100226. doi:10.1016/j.fsisyn.2022.100226

7. Senge T, Madea B, Junge A, Rothschild MA, Schneider PM. STRs, mini STRs and SNPs – A comparative study for typing degraded DNA. Leg Med. 2011 Mar;13(2):68–74. doi:10.1016/j.legalmed.2010.12.001

8. Butler JM. Advanced topics in forensic DNA typing : methodology. Elsevier Academic Press; 2012.

9. Butler J, Shen Y, McCord B. The Development of Reduced Size STR Amplicons as Tools for Analysis of Degraded DNA. J Forensic Sci. 2003 Sep 1;48(5):1–11. doi:10.1520/JFS2003043

10. van Oorschot RA, Ballantyne KN, Mitchell RJ. Forensic trace DNA: a review. Investig Genet. 2010 Dec 1;1(1):14. doi:10.1186/2041-2223-1-14 PubMed PMID: 21122102.

11. Dixon LA, Dobbins AE, Pulker HK, Butler JM, Vallone PM, Coble MD, et al. Analysis of artificially degraded DNA using STRs and SNPs—results of a collaborative European (EDNAP) exercise. Forensic Sci Int. 2006 Dec;164(1):33–44. doi:10.1016/j.forsciint.2005.11.011

12. Grubwieser P, Mühlmann R, Berger B, Niederstätter H, Pavlic M, Parson W. A new “miniSTR-multiplex” displaying reduced amplicon lengths for the analysis of degraded DNA. Int J Legal Med. 2006 Mar 13;120(2):115–20. doi:10.1007/s00414-005-0013-6

13. Kampmann ML, Børsting C, Jepsen AH, Andersen MM, Aagreen CIV, Poggiali B, et al. Preparing for shotgun sequencing in forensic genetics – Evaluation of DNA extraction and library building methods. Forensic Sci Int Genet. 2025 Mar;76:103234. doi:10.1016/j.fsigen.2025.103234

14. Cihlar JC, Woerner AE, King JL, Hawkins JB, Coble MD. Developmental validation of a whole genome sequencing workflow for use in a forensic laboratory. Forensic Sci Int Genet. 2026 Feb;81:103380. doi:10.1016/j.fsigen.2025.103380

15. Green RE, Krause J, Ptak SE, Briggs AW, Ronan MT, Simons JF, et al. Analysis of one million base pairs of Neanderthal DNA. Nature. 2006 Nov 16;444(7117):330–6. doi:10.1038/nature05336

16. Poinar HN, Schwarz C, Qi J, Shapiro B, MacPhee RDE, Buigues B, et al. Metagenomics to Paleogenomics: Large-Scale Sequencing of Mammoth DNA. Science (1979). 2006 Jan 20;311(5759):392–4. doi:10.1126/science.1123360

17. Danielewski M, Żuraszek J, Zielińska A, Herzig KH, Słomski R, Walkowiak J, et al. Methodological Changes in the Field of Paleogenetics. Genes (Basel). 2023 Jan 16;14(1):234. doi:10.3390/genes14010234

18. Zavala EI, Thomas JT, Sturk-Andreaggi K, Daniels-Higginbotham J, Meyers KK, Barrit-Ross S, et al. Ancient DNA Methods Improve Forensic DNA Profiling of Korean War and World War II Unknowns. Genes (Basel). 2022 Jan 11;13(1):129. doi:10.3390/genes13010129

19. Emery MV, Bolhofner K, Winingear S, Oldt R, Montes M, Kanthaswamy S, et al. Reconstructing full and partial STR profiles from severely burned human remains using comparative ancient and forensic DNA extraction techniques. Forensic Sci Int Genet. 2020 May;46:102272. doi:10.1016/j.fsigen.2020.102272

20. Budowle B, Mittelman K, Mittelman D. Genomics will forever reshape forensic science and criminal justice. Genome Biol. 2025 Sep 22;26(1):296. doi:10.1186/s13059-025-03798-x

21. Xie Q, Zhao W, Liu W, Zhao Y, Chen X, Li J, et al. Forensic SNP genealogy inference using whole genome sequencing data of varying depths. Forensic Sci Int Genet. 2025 Sep;79:103296. doi:10.1016/j.fsigen.2025.103296

22. Jepsen AH, Poggiali B, Jensen MT, Kling D, Zavala EI, Børsting C, et al. Preparing for shotgun sequencing in forensic genetics – Benchmarking of tools for read mapping, genotype calling, and imputation. Forensic Sci Int Genet. 2026 Jun;84:103505. doi:10.1016/j.fsigen.2026.103505

23. Jønck CG, Qian X, Simayijiang H, Børsting C. STRinNGS v2.0: Improved tool for analysis and reporting of STR sequencing data. Forensic Sci Int Genet. 2020 Sep;48:102331. doi:10.1016/j.fsigen.2020.102331

24. Brandhagen MD, Loreille O, Irwin JA. Fragmented Nuclear DNA Is the Predominant Genetic Material in Human Hair Shafts. Genes (Basel). 2018 Dec 18;9(12):640. doi:10.3390/genes9120640

25. Loreille O, Tillmar A, Brandhagen MD, Otterstatter L, Irwin JA. Improved DNA Extraction and Illumina Sequencing of DNA Recovered from Aged Rootless Hair Shafts Found in Relics Associated with the Romanov Family. Genes (Basel). 2022 Jan 23;13(2). doi:10.3390/genes13020202 PubMed PMID: 35205247.

26. Larnane A, Lefèvre-Horgues C, Cruaud C, Fund C, Le Floch E, Sandron F, et al. Characterization of challenging forensic DNA traces using advanced molecular technologies. Int J Legal Med. 2025 Jul 1;139(4):1511–27. doi:10.1007/s00414-025-03448-8

27. Gaudio D, Fernandes DM, Schmidt R, Cheronet O, Mazzarelli D, Mattia M, et al. Genome-Wide DNA from Degraded Petrous Bones and the Assessment of Sex and Probable Geographic Origins of Forensic Cases. Sci Rep. 2019 Jun 3;9(1):8226. doi:10.1038/s41598-019-44638-w

28. B⊘rsting C, Mogensen HS, Morling N. Forensic genetic SNP typing of low-template DNA and highly degraded DNA from crime case samples. Forensic Sci Int Genet. 2013 May;7(3):345–52. doi:10.1016/j.fsigen.2013.02.004

29. Sanchez JJ, Phillips C, Børsting C, Balogh K, Bogus M, Fondevila M, et al. A multiplex assay with 52 single nucleotide polymorphisms for human identification. Electrophoresis. 2006 May 27;27(9):1713–24. doi:10.1002/elps.200500671

30. Musgrave-Brown E, Ballard D, Balogh K, Bender K, Berger B, Bogus M, et al. Forensic validation of the SNPforID 52-plex assay. Forensic Sci Int Genet. 2007 Jun;1(2):186–90. doi:10.1016/j.fsigen.2007.01.004

31. Sobrino B, Brión M, Carracedo A. SNPs in forensic genetics: a review on SNP typing methodologies. Forensic Sci Int. 2005 Nov;154(2–3):181–94. doi:10.1016/j.forsciint.2004.10.020

32. Woerner AE, King JL, Budowle B. Fast STR allele identification with STRait Razor 3.0. Forensic Sci Int Genet. 2017 Sep;30:18–23. doi:10.1016/j.fsigen.2017.05.008

33. Mousavi N, Shleizer-Burko S, Yanicky R, Gymrek M. Profiling the genome-wide landscape of tandem repeat expansions. Nucleic Acids Res. 2019 Sep 5;47(15):e90–e90. doi:10.1093/nar/gkz501

34. Willems T, Zielinski D, Yuan J, Gordon A, Gymrek M, Erlich Y. Genome-wide profiling of heritable and de novo STR variations. Nat Methods. 2017 Jun 24;14(6):590–2. doi:10.1038/nmeth.4267

35. National Human Genome Research Institute (NHGRI). https://www.genome.gov/about-genomics/fact-sheets/Sequencing-Human-Genome-cost. 2021. The Cost of Sequencing a Human Genome.

36. Oketch JW, Wain L V., Hollox EJ. A comparison of software for analysis of rare and common short tandem repeat (STR) variation using human genome sequences from clinical and population-based samples. PLoS One. 2024 Apr 1;19(4):e0300545. doi:10.1371/journal.pone.0300545

37. Weisburd B, Tiao G, Rehm HL. Insights from a genome-wide truth set of tandem repeat variation. 2023. doi:10.1101/2023.05.05.539588

38. Han W, Zhang X, Zhang Q, Zhou Z. Next-generation sequencing-based tools or nanopore-based tools: which is more suitable for short tandem repeats genotyping of nanopore sequencing? Bioinformatics advances. 2025;5(1):vbaf119. doi:10.1093/bioadv/vbaf119 PubMed PMID: 40521380.

39. Valle-Silva G, Frontanilla TS, Ayala J, Donadi EA, Simões AL, Castelli EC, et al. Analysis and comparison of the STR genotypes called with HipSTR, STRait Razor and toaSTR by using next generation sequencing data in a Brazilian population sample. Forensic Sci Int Genet. 2022 May;58:102676. doi:10.1016/j.fsigen.2022.102676

40. Ganschow S, Silvery J, Kalinowski J, Tiemann C. toaSTR: A web application for forensic STR genotyping by massively parallel sequencing. Forensic Sci Int Genet. 2018 Nov;37:21–8. doi:10.1016/j.fsigen.2018.07.006

41. Zhang X, Ji X, Wang L, Chi L, Li C, Wen S, et al. STRsensor: a computationally efficient method for STR allele-typing from massively parallel sequencing data. Brief Bioinform. 2024 Nov 22;26(1). doi:10.1093/bib/bbae637

42. Frontanilla TS, Valle-Silva G, Ayala J, Mendes-Junior CT. Open-Access Worldwide Population STR Database Constructed Using High-Coverage Massively Parallel Sequencing Data Obtained from the 1000 Genomes Project. Genes (Basel). 2022 Nov 24;13(12):2205. doi:10.3390/genes13122205

43. Frontanilla TS, de Sousa Ferrari M, Ayala J, Mendes-Junior CT. HipSTR-UI: A cross-platform graphical interface for accessible str genotyping from next-generation sequencing data. Forensic Sci Int Genet. 2026 Jun;84:103456. doi:10.1016/j.fsigen.2026.103456

44. Press MO, Carlson KD, Queitsch C. The overdue promise of short tandem repeat variation for heritability. Trends in Genetics. 2014 Nov;30(11):504–12. doi:10.1016/j.tig.2014.07.008

45. Brookes C, Bright JA, Harbison S, Buckleton J. Characterising stutter in forensic STR multiplexes. Forensic Sci Int Genet. 2012 Jan;6(1):58–63. doi:10.1016/j.fsigen.2011.02.001

46. Woerner A, King J, Budowle B. Flanking Variation Influences Rates of Stutter in Simple Repeats. Genes (Basel). 2017 Nov 17;8(11):329. doi:10.3390/genes8110329

47. Walsh PS, Fildes NJ, Reynolds R. Sequence Analysis and Characterization of Stutter Products at the Tetranucleotide Repeat Locus VWA. Nucleic Acids Res. 1996 Jul 1;24(14):2807–12. doi:10.1093/nar/24.14.2807

48. Bacher JW; Schumm JW; McElfresh KC; Rabbach DR. Pentanucleotide repeats: highly polymorphic genetic markers displaying minimal stutter artifact. 1999.

49. Ellegren H. Microsatellites: simple sequences with complex evolution. Nat Rev Genet. 2004 Jun;5(6):435–45. doi:10.1038/nrg1348

50. Shinde D. Taq DNA polymerase slippage mutation rates measured by PCR and quasi-likelihood analysis: (CA/GT)n and (A/T)n microsatellites. Nucleic Acids Res. 2003 Feb 1;31(3):974–80. doi:10.1093/nar/gkg178

51. Fazekas AJ, Steeves R, Newmaster SG. Improving Sequencing Quality from PCR Products Containing Long Mononucleotide Repeats. Biotechniques. 2010 Apr 3;48(4):277–85. doi:10.2144/000113369

52. Hauge XY, Litt M. A study of the origin of ‘shadow bands’ seen when typing dinucleotide repeat polymorphisms by the PCR. Hum Mol Genet. 1993;2(4):411–5. doi:10.1093/hmg/2.4.411

53. Shi Y, Niu Y, Zhang P, Luo H, Liu S, Zhang S, et al. Characterization of genome-wide STR variation in 6487 human genomes. Nat Commun. 2023 Apr 12;14(1):2092. doi:10.1038/s41467-023-37690-8

54. Kidd KK, Pakstis AJ, Gandotra N, Scharfe C, Podini D. A multipurpose panel of microhaplotypes for use with STR markers in casework. Forensic Sci Int Genet. 2022 Sep;60:102729. doi:10.1016/j.fsigen.2022.102729

55. Oldoni F, Kidd KK, Podini D. Microhaplotypes in forensic genetics. Forensic Sci Int Genet. 2019 Jan;38:54–69. doi:10.1016/j.fsigen.2018.09.009

56. Kidd KK, Pakstis AJ, Speed WC, Grigorenko EL, Kajuna SLB, Karoma NJ, et al. Developing a SNP panel for forensic identification of individuals. Forensic Sci Int. 2006 Dec;164(1):20–32. doi:10.1016/j.forsciint.2005.11.017

57. van der Gaag KJ, de Leeuw RH, Laros JFJ, den Dunnen JT, de Knijff P. Short hypervariable microhaplotypes: A novel set of very short high discriminating power loci without stutter artefacts. Forensic Sci Int Genet. 2018 Jul;35:169–75. doi:10.1016/j.fsigen.2018.05.008

58. Chen P, Yin C, Li Z, Pu Y, Yu Y, Zhao P, et al. Evaluation of the Microhaplotypes panel for DNA mixture analyses. Forensic Sci Int Genet. 2018 Jul;35:149–55. doi:10.1016/j.fsigen.2018.05.003

59. Standage D, Just R, Scharfe C, Gandotra N. Empirical haplotype calling and probabilistic interpretation of microhaplotype profiles. Forensic Sci Int Genet Suppl Ser. 2022 Dec;8:265–7. doi:10.1016/j.fsigss.2022.10.057

60. Jønck CG, Børsting C. Introduction of the python script MHinNGS for analysis of microhaplotypes. Forensic Sci Int Genet Suppl Ser. 2022 Dec;8:79–81. doi:10.1016/j.fsigss.2022.09.029

61. Zhang C, Cao YD, Song JJ, Rao M, Nie SJ, Zhang GF, et al. MHTyper: a microhaplotype allele-calling pipeline for use with next generation sequencing data. Australian Journal of Forensic Sciences. 2021 May 4;53(3):283–90. doi:10.1080/00450618.2019.1699956

62. Ji X, Feng Y, Chi L, Fu M, Kang K, Zhang C, et al. SMART-MHmix: A probabilistic model for microhaplotype-based forensic DNA mixture analysis. Forensic Sci Int Genet. 2026 Jun;84:103509. doi:10.1016/j.fsigen.2026.103509

63. Verogen. https://www.qiagen.com/dk/products/human-id-and-forensics/nextgeneration-sequencing/verogen-forenseq-mainstay-kit?catno=V16000183. ForenSeq MainstAY SE Kit.

64. R Core Team. R: A Language and Environment for Statistical Computing [Internet]. Vienna, Austria; 2021. Available from: https://www.R-project.org/

65. Wickham H. ggplot2: Elegant Graphics for Data Analysis [Internet]. Springer-Verlag New York; 2016. Available from: https://ggplot2.tidyverse.org

66. 1000 Genomes Project. https://42basepairs.com/browse/s3/1000genomes/1000G_2504_high_coverage.

67. Byrska-Bishop M, Evani US, Zhao X, Basile AO, Abel HJ, Regier AA, et al. High-coverage whole-genome sequencing of the expanded 1000 Genomes Project cohort including 602 trios. Cell. 2022 Sep 1;185(18):3426–3440.e19. doi:10.1016/j.cell.2022.08.004 PubMed PMID: 36055201.

68. https://github.com/HipSTR-Tool/HipSTR-references/tree/master/human.hg38.hipstr_reference.bed.gz.

69. Quinlan AR, Hall IM. BEDTools: a flexible suite of utilities for comparing genomic features. Bioinformatics. 2010 Mar 15;26(6):841–2. doi:10.1093/bioinformatics/btq033

70. Smedley D, Haider S, Ballester B, Holland R, London D, Thorisson G, et al. BioMart – biological queries made easy. BMC Genomics. 2009 Dec 14;10(1):22. doi:10.1186/1471-2164-10-22

71. Harrison PW, Amode MR, Austine-Orimoloye O, Azov AG, Barba M, Barnes I, et al. Ensembl 2024. Nucleic Acids Res. 2024 Jan 5;52(D1):D891–9. doi:10.1093/nar/gkad1049

72. Karolchik D. The UCSC Table Browser data retrieval tool. Nucleic Acids Res. 2004 Jan 1;32(90001):493D – 496. doi:10.1093/nar/gkh103

73. https://emea.illumina.com/products/by-type/informatics-products/designstudio.html. DesignStudio Assay Design Tool.

74. Pedersen MW, Antunes C, De Cahsan B, Moreno-Mayar JV, Sikora M, Vinner L, et al. Ancient Human Genomes and Environmental DNA from the Cement Attaching 2,000-Year-Old Head Lice Nits. Mol Biol Evol. 2022 Feb 3;39(2). doi:10.1093/molbev/msab351

